# HuBMAPR: an R Client for the HuBMAP Data Portal

**DOI:** 10.1101/2024.09.26.615227

**Authors:** Christine Hou, Shila Ghazanfar, Federico Marini, Martin Morgan, Stephanie C. Hicks

**Author notes:** Corresponding author. Department of Biostatistics, Johns Hopkins Bloomberg School of Public Health, 615 N Wolfe St, Baltimore, MD, 21205, United States.

## Abstract

**Summary:** The Human BioMolecular Atlas Program (HuBMAP) constructs the worldwide available platform to research the human body at the cellular level. The HuBMAP Data Portal encompasses a wide range of data resources measured on emerging experimental technologies at a spatial resolution. To broaden access to the HuBMAP Data Portal, we introduce an R client called `HuBMAPR` available on Bioconductor. This gives an efficient and programmatic interface, enabling researchers to discover and retrieve HuBMAP data easier and faster.

**Availability:** This package is available on GitHub (https://github.com/christinehou11/HuBMAPR) and has been submitted to Bioconductor.

## 1 Introduction

The Human BioMolecular Atlas Program (HuBMAP) is a comprehensive, open-sourced, global atlas of the human body at a cellar resolution (HuBMAP Consortium, 2019; Jain et al., 2023). With the development of computational tools, researchers aim to understand the interaction, spatial organization, and specialization of the estimated trillions of cells in the adult human body contribute to organ and tissue function, which further helps to understand their relationship with the human health. Data are hosted on the HuBMAP Data Portal^1^ and permitted contributors can upload experimental data to the HuBMAP Data Portal following the data submission guide^2^.

HuBMAP data generally encompasses five primary entity categories: (i) dataset, (ii) donor, (iii) sample, (iv) collection, and (v) publication. As of August 2024, more than 2,300 datasets are available across 20 assay types, including Co-detection by indexing (CODEX)^3^, Imaging Mass Cytometry (IMC)^4^, bulk and single-cell RNA Sequencing (RNAseq)^5^, and Sequential Fluorescence In-Situ Hybridization (seqFISH)^6^ (**Fig. S1**), from 228 donors and 1,928 samples across 31 different organs. HuBMAP datasets generated from related experiments (or sharing similar characteristics) are grouped into 18 HuBMAP collections^7^. Despite these data resources being available on the HuBMAP Data Portal, there currently lacks programmatic interface in R (R Core Team, 2024) to access, explore, retrieve, and download these data. In this work, we address the problem by developing the `HuBMAPR` R/Bioconductor (Robert et al., 2004) package to enable researchers to programmatically explore and download HuBMAP data.

## 2 HuBMAPR **R/Bioconductor package**

### 2.1 Overview of HuBMAP Data Portal and HuBMAPR

The `HuBMAPR` package retrieves data from the same five entity categories in HuBMAP using three different identifiers: (i) HuBMAP ID, (ii) Universally Unique Identifier (UUID), and (iii) Digital Object Identifiers (DOI). The HuBMAPR package primarily uses the UUID–a 32-digit hexadecimal number–and the more human-readable HuBMAP ID (**Fig. 1B**). Considering precision and compatibility with software implementation and data storage, UUID serves as the primary identifier to retrieve data across various functions, with the UUID mapping uniquely to its corresponding `HuBMAP` ID. A systematic nomenclature is adopted for functions in the package by appending the entity category prefix to the concise description of the specific functionality. For example, `dataset detail()` helps to retrieve the detailed metadata of one specific dataset, and `donor derived()` provides the derived datasets of specific donors. Most of the functions are grouped by entity categories, simplifying the process of selecting the appropriate functions to retrieve desired information associated with a UUID from the specific entity category. The structure of these functions is consistent across all entity categories, with some minor exceptions for collection and publication entities.

**Fig 1.**
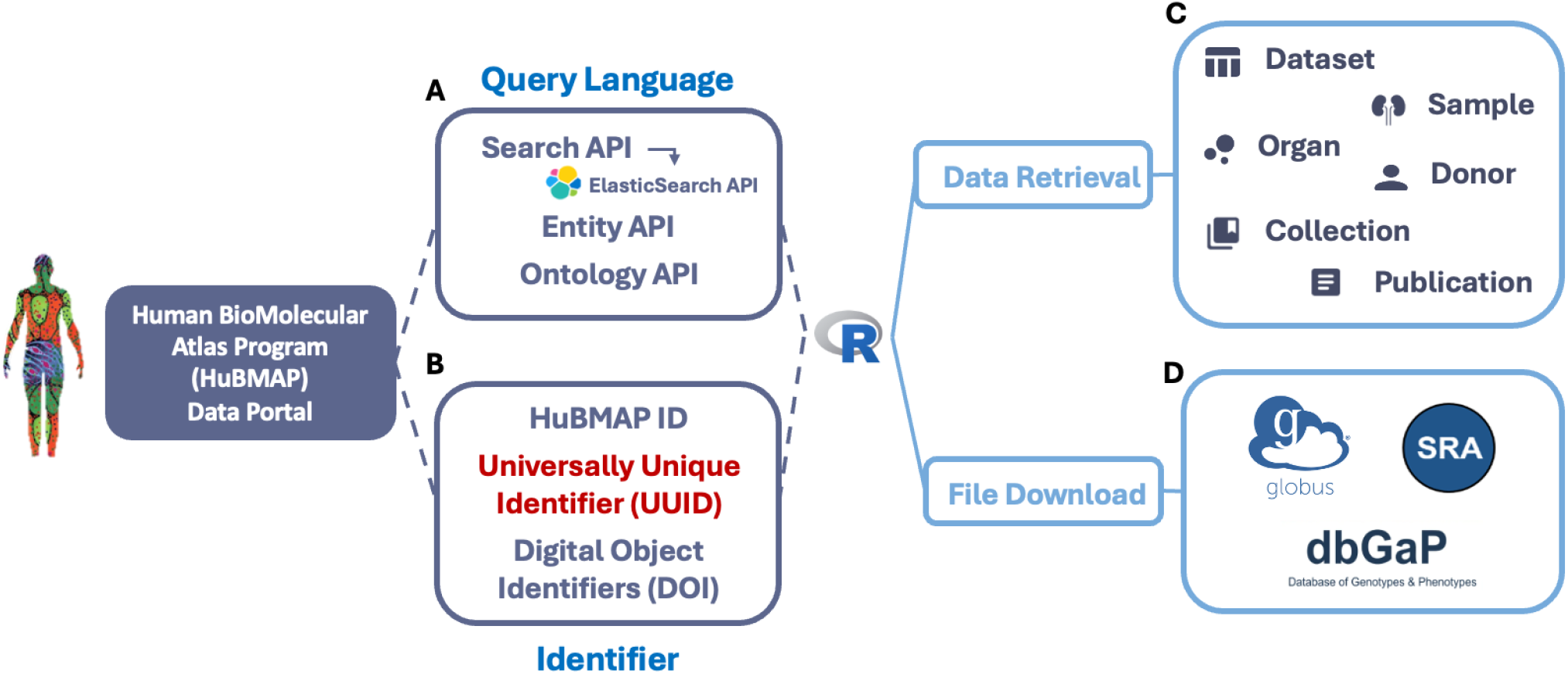
`HuBMAPR` builds a programmatic interface between the HuBMAP data portal and the R programming language utilizing multiple APIs based on (**A**) a query language and (**B**) extracting specific data entry based primarily on Universally Unique Identifiers (UUIDs). (**C**) Within R, the `HuBMAPR` package helps to explore and to retrieve data from different entity categories. (**D**) Data files can be accessed and downloaded via Globus, NCBI Database of Genotypes and Phenotypes (dbGAP), or Sequence Read Archive (SRA).

### 2.2 Data retrieval

The `HuBMAPR` package arranges HuBMAP data chronologically by the last modification date, providing extensive donor physical, social, or ethnic demographic characteristics including biological sex at birth, age, self-reported race, organ, body mass index, and other metadata. Additionally, other features include experimental statistics such as analyte class, processing pipeline^8^, sample category, affiliation information (e.g., contributor contacts, attribution group name, registration institution), and status updates such as publication date, publication status, and last modification date. By carefully selecting and presenting these data details from the HuBMAP Data Portal, the `HuBMAPR` package offers a robust R-based interface for comprehensive data discovery, filtering, and extraction from datasets, samples, and donors. Moreover, some additional functions are available to retrieve the summary of organs and relevant datasets and textual descriptions of collections and publications (**Fig. 1C**).

In addition to generating details of entity data information in a structured and accessible format, the package facilitates the retrieval of data provenance, encompassing both ancestors and descendants. HuBMAP and the `HuBMAPR` package defines an ancestor record as an individual record from which a specific donor, sample, or dataset is derived. Conversely, a descendant record is defined as an individual record that has been derived from other preceding records. The donor initiates the provenance hierarchy, with the donor and the donor-derived sample organ regarded as foundational elements. Specific sample categories, such as section, suspension, and block, can be derived from the sample organ. Various assay types are applied to generate HuBMAP datasets from the samples. The provenance hierarchy culminates in the supporting dataset, particularly when the dataset is further processed by specific pipelines such as snapATAC (Rongxin et al., 2021), Salmon (Patro et al., 2017), or Cytokit (Eric et al., 2019). Corresponding functions are available to retrieve ancestor and descendant records for the donor, sample, and dataset. Due to the definition of the collection and publication entity, there is no function to retrieve ancestors and descendants.

### 2.3 Files delivery

The HuBMAP Data Portal offers access to both open and restricted access to individual record files, adhering to the NIH Genomics Data Sharing (GDH) Policy and other relevant legal frameworks^9^. The public HuBMAP data can be accessed via Globus^10^, a secure and efficient cloud platform for large-size data storage and rapid file transfers (**Fig. 1D**). Using the unique dataset UUID, the `HuBMAPR` package connects to the HuBMAP public collection within the Globus research data management system, directing the users to the Globus online website to preview and download raw data products, downstream analysis reports, metadata files, and visualizations. We have developed an experimental R package named `rglobus` (Morgan, 2024) to discover and navigate Globus collections and to transfer files and directories between collections. Additional `rglobus` functionality is under development.

Restricted-access databases contain human-protected sequencing data, requiring special permissions. The NIH Data Access Committee manages access to restricted databases, and users must authenticate their identities to request downloads. Except for HuBMAP Consortium members, access to restricted-access databases may be granted through The database of Genotypes and Phenotypes (dbGaP) (Tryka et al., 2014) or Sequence Read Archive (SRA) (Katz et al., 2022), if available. However, it is possible that neither dbGaP nor SRA can provide direct download links, and these datasets may take additional time and effort before they become accessible. Therefore, if the requested data files are restricted, `HuBMAPR` package will return either helpful links (dbGAP or SRA) or messages to provide clarifications and instructions.

## 3 Conclusion

The HuBMAPR package offers programmatic and efficient interface to access and retrieve HuBMAP data in R. By building a local platform, the package helps to explore HuBMAP data within R leveraging the `tidyverse` (Wickham et al., 2019) for further data wrangling. The `HuBMAPR` package returns detailed data summaries from different entity categories in an organized format, and connects to the external Globus data management system for file browsing and download. Downloaded data can then be explored in R and Bioconductor packages and other software.

## 4 Supplementary Data

Supplementary Data is available at Bioinformatics online.

## 5 Author contributions

C.H.: Conceptualization, Methodology, Software, Validation, Formal analysis, Investigation, Data Curation, Writing, Visualization; M.M: Conceptualization, Software, Code Review; D.M: Conceptualization, Code Review; S.G.: Code Review; S.C.H.: Conceptualization, Resources, Writing Review & Editing, Supervision, Project administration, Funding acquisition. All co-authors approved the final manuscript.

## 8 Data Availability

All published data is available on the HuBMAP Data Portal^11^. The open-source software package `HuBMAPR` is based on R programming language (R Core Team, 2024) and the software license we are using is Artistic 2.0^12^. This package is available on GitHub (https://github.com/christinehou11/HuBMAPR) and has been submitted to Bioconductor.

## Supplementary Materials

**Supplementary Figure S1.**
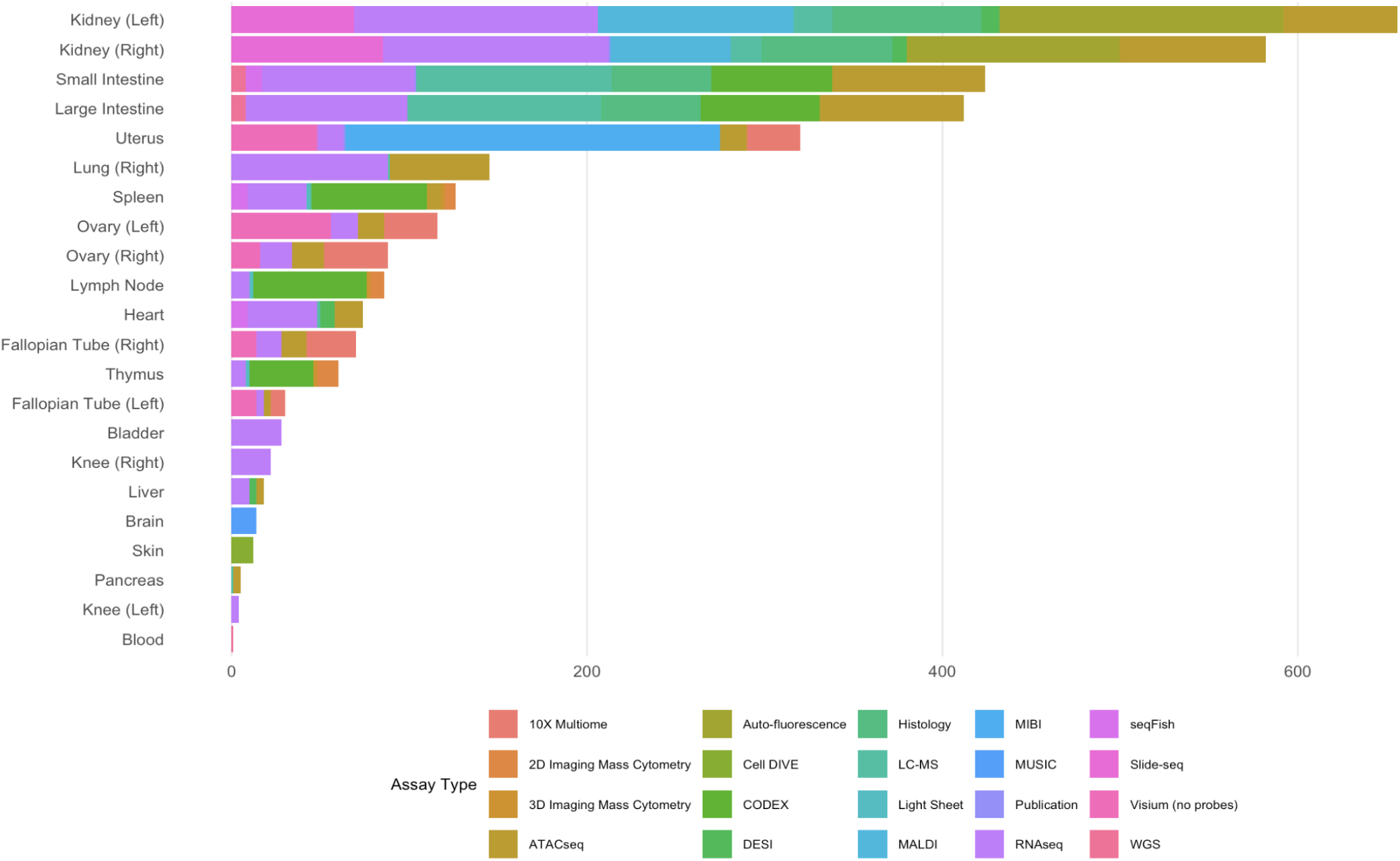
Overview of tissue types in HuBMAP. HuBMAP Data Portal consists of the original datasets and corresponding support/processed datasets collected from a wide range of organs and processed by multiple assay types (as of August 2024)

## Footnotes

1 https://portal.hubmapconsortium.org/

2 https://docs.hubmapconsortium.org/data-submission

3 https://docs.hubmapconsortium.org/assays/codex

4 https://docs.hubmapconsortium.org/assays/imc

5 https://docs.hubmapconsortium.org/assays/rnaseq

6 https://docs.hubmapconsortium.org/assays/seqfish

7 https://portal.hubmapconsortium.org/collections

8 https://github.com/hubmapconsortium

9 https://hubmapconsortium.org/policies/

10 https://www.globus.org/

11 https://portal.hubmapconsortium.org/

12 https://www.perlfoundation.org/artistic-license-20.html

